# Molecular layer disinhibition unlocks climbing-fiber-instructed motor learning in the cerebellum

**DOI:** 10.1101/2023.08.04.552059

**Authors:** Ke Zhang, Zhen Yang, Michael A. Gaffield, Garrett G. Gross, Don B. Arnold, Jason M. Christie

## Abstract

Climbing fibers supervise cerebellar learning by providing signals to Purkinje cells (PCs) that instruct adaptive changes to mistakenly performed movements. Yet, climbing fibers are regularly active, even during well performed movements, suggesting that a mechanism dynamically regulates the ability of climbing fibers to induce corrective plasticity in response to motor errors. We found that molecular layer interneurons (MLIs), whose inhibition of PCs powerfully opposes climbing-fiber-mediated excitation, serve this function. Optogenetically suppressing the activity of floccular MLIs in mice during the vestibulo-ocular reflex (VOR) induces a learned increase in gain despite the absence of performance errors. Suppressing MLIs when the VOR is mistakenly underperformed reveled that their inhibitory output is necessary to orchestrate gain-increase learning by conditionally permitting climbing fibers to instruct plasticity induction during ipsiversive head turns. Ablation of an MLI circuit for PC disinhibition prevents gain-increase learning during VOR performance errors which was rescued by re-imposing PC disinhibition through MLI activity suppression. Our findings point to a decisive role for MLIs in gating climbing-fiber-mediated learning through their context-dependent inhibition of PCs.

## Introduction

The accuracy and efficiency of movement is improved by adaptive learning triggered in response to mistakes (when motor intent and outcome conflict). However, in the absence of motor errors (when the outcome matches intent), adaptation may have little benefit to behavioral outcomes because the movements were already well performed. Therefore, neural circuits mediating supervised motor learning must be computationally flexible such that corrective plasticity is principally induced when adaptation is necessary, for example, in response to a motor error, but are otherwise resistant to change. In the cerebellar cortex, a key brain center for implicit motor learning, climbing fibers fire during motor errors reliably eliciting signals in postsynaptic Purkinje cells (PCs) that instruct corrective plasticity induction (Llinas and Sugimori, 1980; Kitazawa et al., 1998; Ozden et al., 2009). However, climbing fibers also discharge during properly performed behaviors (Heffley et al., 2018; Kostadinov et al., 2019). Therefore, additional mechanisms, beyond the activity of climbing fibers alone, must be engaged to fully explain the neural circuit basis of supervised cerebellar learning (Knudsen, 1994; Raymond and Medina, 2018).

PCs also integrate activity from GABA-releasing molecular layer interneurons (MLIs). Inhibition from MLIs curtails climbing-fiber-mediated excitation of PCs (Brunel et al., 2004; Mittmann et al., 2005; Dizon and Khodakhah, 2011; Kitamura and Hausser, 2011). By opposing climbing-fiber-evoked signaling, MLI-to-PC inhibition can gate the induction of long-term depression (LTD) at parallel fiber synapses, a plasticity type believed to be important for many forms of cerebellar learning (Callaway et al., 1995; Gaffield et al., 2018; Rowan et al., 2018). As such, MLI activity could prevent climbing fiber signaling from inducing unproductive plasticity during behavioral contexts that do not warrant adaptive change. Gating implies that there must be a mechanism to unlock the ability of climbing fibers to induce plasticity during erroneous motor contexts that require corrective learning. In the forebrain, a structured interconnectivity among local interneuron networks promotes pyramidal cell disinhibition, thus forming a neural circuit mechanism that, when actively engaged in a context-dependent manner, alleviates inhibitory constraints over plasticity and learning (Kepecs and Fishell, 2014; Letzkus et al., 2015). Because MLIs in the cerebellar cortex are inhibited by other MLIs, PCs, and PC-layer interneurons, these connections form a circuit for disinhibiting PCs (O’Donoghue et al., 1989; Blot et al., 2016; Witter et al., 2016; Arlt and Häusser, 2020; Osorno et al., 2021; Halverson et al., 2022). Whether PC disinhibitory circuits play a role in regulating error-driven cerebellar learning is currently unknown.

In this current study, we investigated the circuit mechanisms in the cerebellar cortex that underlie gated cerebellar learning. Using neural activity measurements and manipulations in combination with behavioral analysis, we identified the importance of MLI-mediated inhibition of PCs in preventing adaptation in mistake-free contexts and the necessity of an MLI-mediated circuit for PC disinhibition that conditionally permits climbing fibers to instruct motor adaptation in response to performance errors. These discoveries amend rules governing cerebellar function, indicating that supervised learning requires the coordinated interplay of activity within the MLI ensemble to conditionally gate the effectiveness climbing fibers to instruct corrective plasticity in response to motor errors.

## Results

### Context-dependent adaptation of VOR gain

We investigated the neural circuit mechanisms underlying the conditional nature of cerebellar learning using the vestibulo-ocular reflex (VOR) as a behavioral readout in mice. The VOR helps stabilize images on the retina by triggering compensatory eye movements in response to vestibular stimuli elicited by head turns. Image motion across the eye (retinal slip) indicates VOR performance errors (Robinson, 2022). The direction of retinal slip relative to that of the head turn indicates whether the eye movements were too small or too large to compensate for the self-motion. The presence of retinal slip during the VOR triggers corrective leaning, instantiated in the cerebellum, to increase or decrease the response gain (amplitude). This learning restores accurate performance (Ito, 1982). We induced the sensory perception of either an under- or over-performing VOR in head-fixed mice by associatively pairing horizontal sinusoidal head turns with a visual stimulus that moved either in the opposite- or same-direction of the head, respectively (**Supplemental Figure 1A,C**). Training sessions consisting of the repeated pairing of either stimulus condition (60 min) drove an expected adaptive increase or decrease in VOR gain for visual and head motion stimuli that moved in the opposite or same direction, respectively, relative to a baseline measurement obtained immediately prior to training (**Supplemental Figure 1B,D**).

Reflexive movements, such as the VOR, should ideally be resistant to changes in the absence of motor errors, as inadvertent modifications could degrade subsequent performance. To test if the VOR gain is resistant to alteration when elicited in a context with little retinal slip, we paired head turns with a stationary visual stimulus (**Figure 1A**). The lack of an induced gain change during training (**Figure 1B**) indicated that this condition was non-adaptive, in stark contrast to the effect of opposite- or same-direction head and visual motion on the VOR.

**Figure 1.**
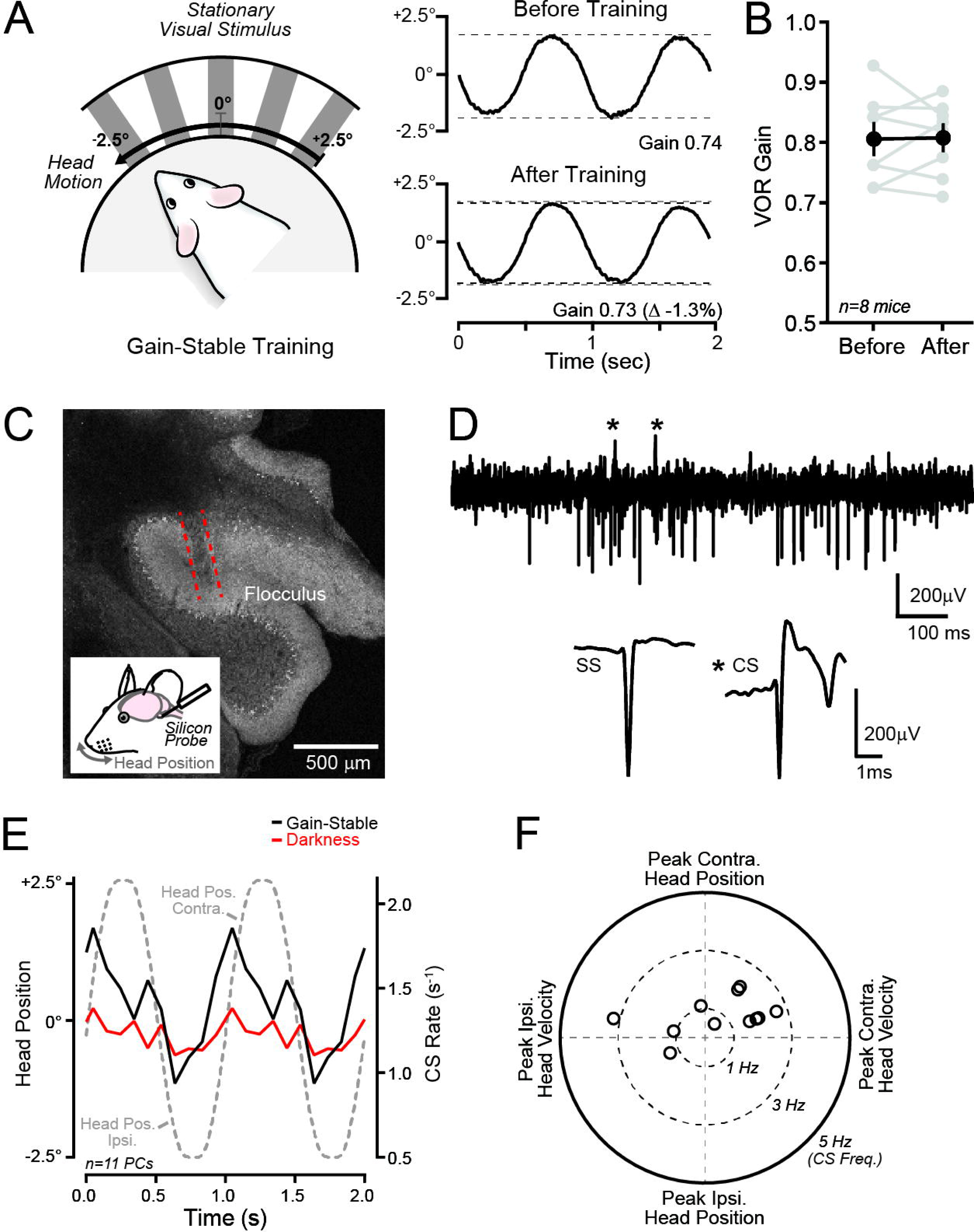
PC complex spiking in a minimally adapting VOR context. A. Left panel: Gain-stable training stimuli involved sinusoidal head turns (1 Hz; ±5.0°) paired with a stationary visual stimulus (60 min). Right panel: Average VOR-evoked eye movements, isolated in darkness, from a mouse obtained before and after training. The light gray dotted line on the bottom eye trace indicates the gain measurement obtained immediately prior to training. B. VOR performance was unaffected by the gain-stable training procedure across mice. Measurements from individual mice are denoted in gray with mean ± SEM in black (p = 0.8954; paired t-test). C. Extracellular electrophysiological recordings were obtained from the flocculus. The histological image shows the post-recording lesion from an example mouse. D. Recording from a PC with example simple spikes (SS) and complex spikes (CS) highlighted below (complex spikes marked with asterisks). E. Average complex spike activity in the same floccular PCs during sinusoidal head turns in darkness or with a stationary visual surround (i.e., the gain-stable training stimulus; n = 5 mice). F. Polar plot showing the preferred tuning of complex spiking in individual floccular PCs during the gain-stable training stimulus.

### Modulation of Purkinje cell complex spiking during a non-adaptive VOR context

VOR learning is known to involve the flocculus, which is a specialized region of the vestibulo-cerebellum (Ito, 1982). In this region, visual stimuli modulate climbing-fiber-induced complex spiking in PCs (Simpson and Alley, 1974; Ghelarducci et al., 1975; Graf et al., 1988; Stone and Lisberger, 1990). During the VOR, climbing fibers reporting retinal slip evoke signals that instruct plasticity induction underlying learned increases to the response gain (Boyden et al., 2006; Kimpo et al., 2014). To determine whether the absence of VOR gain adaptation in the context of a stationary visual stimulus was due to the lack climbing-fiber-mediated activity modulation, we conducted extracellular electrophysiology recordings from PCs *in vivo*. We used high-electrode-density silicon probes that were targeted to the left flocculus (**Figure 1C**) and identified complex spikes in algorithmically sorted PC units by their characteristic waveform (**Figure 1D**).

Stimulating the VOR by passive sinusoidal head turns in darkness produced little change in complex spiking relative to the spontaneous rate measured in quiescence (mean rate: 1.24 ± 0.09 Hz and 1.29 ± 0.07 Hz, respectively; p = 0.40; paired t test, n =11 PCs from 6 mice). By contrast, when the VOR was elicited in the presence of a stationary visual stimulus, the rate of PC complex spiking increased near the peak velocity phases of both ipsiversive (leftward) and contraversive (rightward) head turns (**Figure 1E**). Individual PCs exhibited opposing patterns of climbing-fiber-mediated activity. In many PCs (7/11), the complex spiking rate increased during contraversive (rightward) head turns, while in the remaining cells, the rate of complex spiking increased during ipsiversive (leftward) head turns (**Figure 1F**). Therefore, although floccular PCs showed robust complex spike modulation when the VOR was elicited during a stationary visual stimulus, this activity appeared to be ineffective at inducing gain adaptation. These results suggest that circuit-level processes must gate the effectiveness of climbing fibers to induce cerebellar learning in a context-dependent manner.

### MLI activity suppression induces gain-adaptation in the absence of performance errors

Prior studies have shown that optogenetically activating floccular MLIs prevents gain-increase VOR learning because the resulting inhibition of PCs opposes the climbing-fiber-mediated signals that instruct corrective plasticity induction (Callaway et al., 1995; Kitamura and Hausser, 2011; Gaffield et al., 2018; Rowan et al., 2018). Therefore, we hypothesized that if MLIs normally inhibit PCs when the VOR generates fully compensatory eye movements in the lighted environment, then MLI-mediated activity could countermand the ability of visually evoked climbing-fiber-mediated signals in PCs to induce plasticity that would alter an otherwise well-calibrated VOR. To test this idea, we used optogenetics to suppress MLI activity while the VOR was repeatedly paired with a stationary visual stimulus.

MLIs in both the left and right flocculus of *Kit::Cre* mice (Amat et al., 2017) were transduced with the inhibitory opsin eNpHR3.0 using a Cre-dependent AAV and light was delivered to the injection sites through implanted optical fibers (**Figure 2A**). In different sessions of training, light pulses for optogenetic actuation were timed to the peak velocity phase of either ipsiversive head turns (i.e., for the left and right flocculus, during leftward and rightward motion, respectively; **Figure 2B,C**) or contraversive head turns (i.e., for the left and right flocculus, during rightward and leftward motion, respectively), epochs during which complex spike firing increased in PCs. In an separate set of eNpHR3.0-expressing *Kit::Cre* mice in which floccular PCs were additionally transduced with GCaMP6f (**Supplemental Figure 2A,B**) for photometry measurements, the MLI optogenetic perturbation resulted in increased PC population activity during the gain-stable VOR stimulus (**Supplemental Figure 2C-E**), indicative of a strong disinhibitory effect.

**Figure 2:**
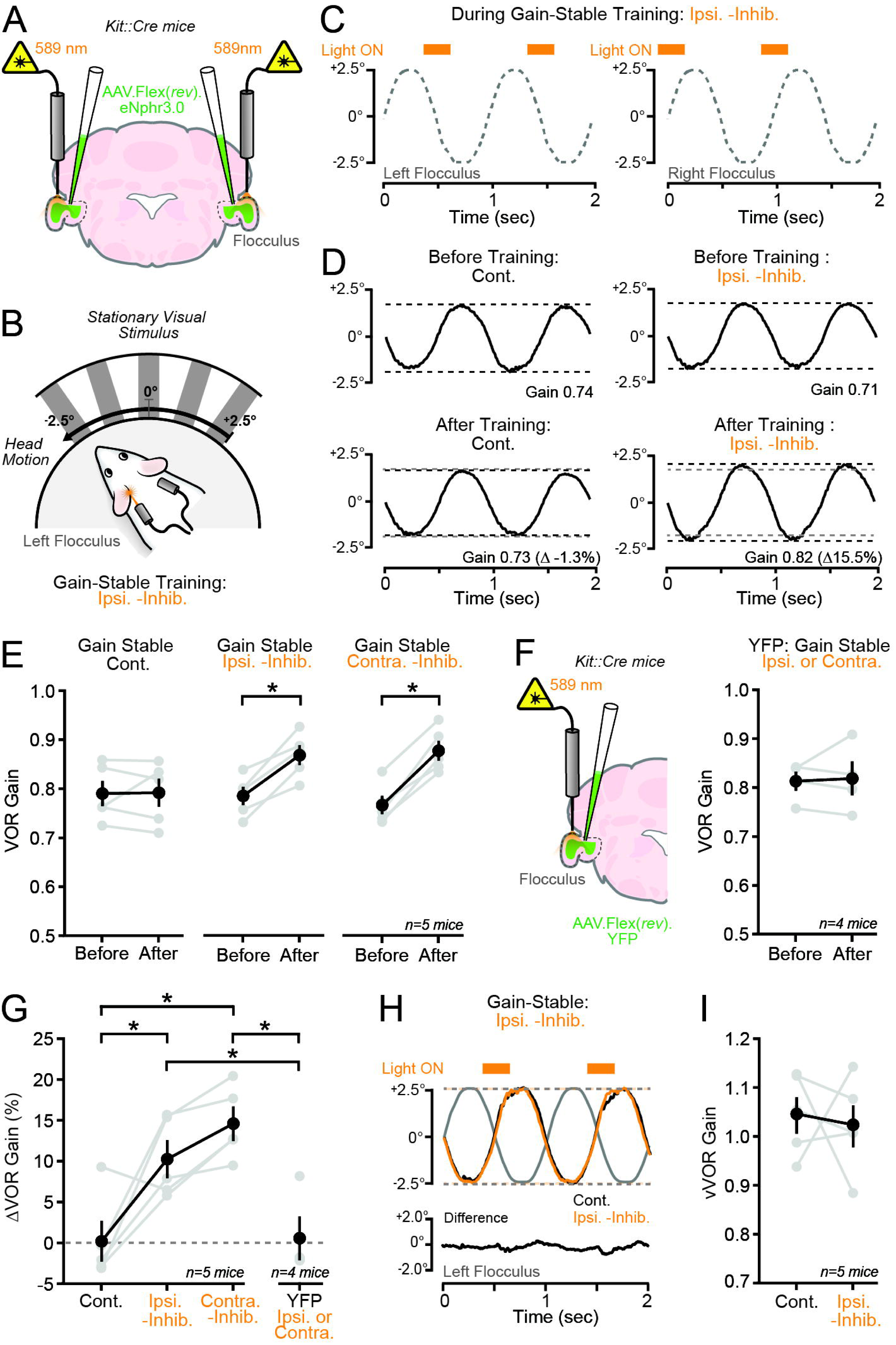
Impact of MLI activity suppression on VOR performance in a minimally adapting context. A. Bilateral injection of Cre-dependent eNphr3.0 AAV in *Kit::Cre* mice, followed by optical fiber implantation, to enable optogenetic suppression of transduced MLIs in both flocculi. B,C. Gain-stable training sessions involved suppressing MLI activity either during ipsiversive head turns (e.g. for the left flocculus during leftward head turns, as illustrated in panel B) or contraversive head turns (60 min). Panel C depicts the timing of the light pulses (λ = 589 nm; 8 mW; 240 ms) during the peak velocity phase of ipsiversive head turns with respect to the left and right flocculus. D. Average VOR-evoked eye movements from the same mouse in different training sessions. Left panel: Measurements from a control session without optogenetic perturbation, taken immediately before and after training. Right panel: In a separate session with MLI activity suppression during ipsiversive head turns, conducted several days later. E. Summary of the influence of MLI activity suppression during gain-stable training on VOR performance across the different conditions in the same set of mice (left to right plots: p = >0.99, *0.0024, and *0.0003; ANOVA with Bonferroni’s post-tests). F. Control mice expressing YFP in place of eNphr3.0 (bilateral injections not illustrated) were trained under the same conditions (p = 0. 0.8137, paired t-test). G. Data from panel E presented as the change in gain (ΔVOR) relative to the baseline response obtained at the beginning of each testing session for each mouse (Cont. vs Ipsi. -Inhib., p = *0.0207; Cont. vs Contra. -Inhib., p = *0.0018; Cont. vs YFP, p = 0.9988; Ipsi. -Inhib. vs Contra. -Inhib., p = 0.4547; Ipsi. -Inhib. vs YFP, p = *0.0252; Contra. -Inhib. vs YFP p = *0.0025; ANOVA with Tukey’s post-tests). H,I. Average visually assisted (v)VOR-evoked eye movements in an eNphr3.0-expressing mouse during pairing with a stationary visual surround, either in the control condition or with optogenetic suppression of MLIs. Both trial types were obtained in rapid succession at the session start. Summary plots compare VOR gain between the two conditions across mice (p = 0.7708, paired t-test). In plots, individual mice are denoted in gray with mean ± SEM; asterisks indicate significance.

For both training conditions, suppressing MLI activity during the pairing procedure led to a marked increase in VOR gain (**Figure 2D-E,G**). Control training sessions in the same mice without the optogenetic perturbation confirmed that the VOR gain remained unchanged (**Figure 2D-E,G**). The specificity of the optogenetic effect to elicit adaptation was demonstrated by the lack of alteration in VOR gain during training when light pulses were delivered to the flocculi of non-eNpHR3.0-expressing *Kit*::*Cre* mice (**Figure 2F,G**). The gain of VOR-evoked eye movements during initial pairing trials that included MLI activity suppression were no different than baseline trials without the optogenetic perturbation (**Figure 2H,I)**. Because eye movements were unimpaired by the optogenetic perturbation, this result rules out the possibility that the gain-increase learning resulted from optogenetically induced VOR performance errors. Notably, the direction and magnitude of the acquired change in VOR gain in the MLI suppression conditions were similar to that produced by opposite-direction visual-head motion mismatches (see **Supplemental Figure 1B**), a context in which climbing fibers instruct the learned gain-increase change (Raymond and Lisberger, 1998; Kimpo et al., 2014). We conclude that MLI activity suppression in a normally non-adaptive context leads to learned change to the VOR gain. Provided that the mice had a well-calibrated VOR prior to training, the learned alteration induced by MLI activity suppression was maladaptive.

### Context-dependent regulation of error-driven motor learning by MLIs

We next tested whether MLI activity suppression influences error driven VOR learning. Mice expressing eNpHR3.0 in floccular MLIs were trained in sessions that included gain-increase stimuli (opposite-direction visual-head motion mismatches; **Figure 3A**). In this context, climbing fibers modulate their firing to the resulting retinal slip. However, climbing-fiber-mediated signaling is only effective at instructing gain-increase learning during ipsiversive, but not contraversive head turns suggesting a permissibility gate, tuned to the direction of head motion, for selective plasticity induction in PCs (Kimpo et al., 2014; Bonnan et al., 2021). We found that when the light pulses for MLI optogenetic suppression were timed to the peak velocity phase of ipsiversive head turns, the mice adapted their VOR by increasing the gain of the response (**Figure 3B**). The direction and magnitude of learning was comparable to that observed in the same mice during control training sessions consisting of pairing opposite-direction visual-vestibular motion mismatches alone (**Figure 3B,C**). The lack of an effect on learning implies that MLIs do not oppose climbing-fiber-mediated instructive signaling during this phase vestibular stimulation. By contrast, when the light pulses were timed to the peak velocity phase of contraversive head turns, the VOR gain remained unchanged relative to the baseline response measured immediately prior to training (**Figure 3B,C**). Thus, even though the mice experienced retinal slip, MLI activity suppression during contraversive head turns obviated the acquisition of an adaptive change to the VOR performance.

**Figure 3:**
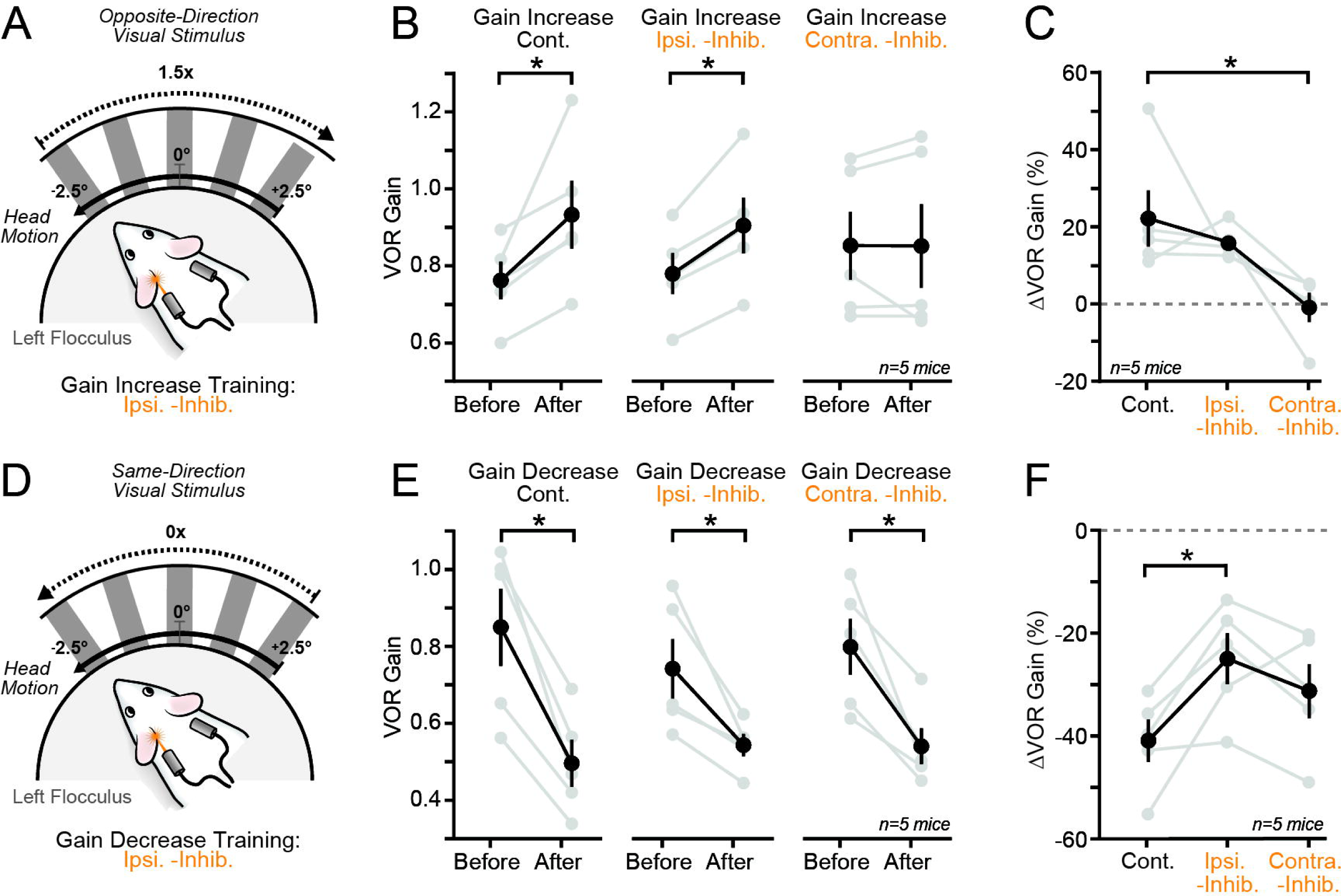
Impact of MLI activity suppression on VOR performance in adapting contexts. A. In training sessions involving the gain-increase stimulus (60 min), floccular MLIs of eNphr3.0-expressing *Kit::Cre* mice were suppressed during the peak velocity phase of either ipsiversive (illustrated for the left flocculus) or contraversive heads turns (λ = 589 nm; 8 - 10 mW; 240 ms). B. Summary data of the effect of MLI activity suppression on the VOR gain of the same set of mice receiving gain-increase training across conditions (left to right plots: p = *0.0092, *0.0468, and >0.99; ANOVA with Bonferroni’s post-tests). C. Data from panel B presented as the change in gain (ΔVOR) relative to the baseline response obtained for each mouse at the start of a session (Cont. vs Ipsi. -Inhib., p = 0.6357; Cont. vs Contra. -Inhib., p = *0.0245; and Contra. vs Ipsi. -Inhib., p = 0.0960; ANOVA with Tukey’s post-tests). D. Depiction of the gain-decrease training procedure during which MLI activity was suppressed during either ipsiversive (illustrated for the left flocculus) or contraversive head turns. E,F. As in panels B and C, but for the gain-decrease training sessions (panel E, left to right plots: p = *0.0001, *0.0053, and *0.0010; ANOVA with Bonferroni’s post-tests; panel F: Cont vs Ipsi. -Inhib, p = *0.0344; Cont vs Contra. -Inhib, p = 0.2039; and Contra. -Inhib vs Ipsi. -Inhib, p = 0.4721; ANOVA with Tukey post-tests). In plots, individual mice are denoted in gray with mean ± SEM; asterisks indicate significance.

We also investigated whether MLI activity suppression affected VOR gain-decrease adaptation (**Figure 3D**). Unlike gain-increase adaptation, this type of VOR learning relies on a plasticity mechanism that does not require instructive signals from climbing fibers (Boyden et al., 2004; Kimpo et al., 2014). eNpHR3.0-expressing mice acquired a learned reduction in VOR gain during training sessions that included light pulses timed to either ipsiversive or contraversive head turns (**Figure 3E,F**). However, for training sessions with light pulses timed to ipsiversive head turns, the magnitude of the learned reduction in VOR gain was reduced compared to control training sessions consisting of gain-decrease stimuli (same-direction visual-head motion mismatches) without the optogenetic perturbation that were obtained in the same mice (**Figure 3F**). We conclude that gain-decrease VOR adaptation is weakened by MLI activity suppression timed to ipsiversive head turns.

### Context-dependent regulation of climbing-fiber-mediated learning by MLIs

A possible explanation for the disparate, conditional effect of MLI activity suppression on gain-increase learning is that, in the absence of a head-direction-selective plasticity gating mechanism mediated by MLIs, climbing fibers instruct performance changes to both phases of retinal slip resulting in an isometric effect on VOR gain. Therefore, to causally test whether MLI activity suppression unlocks the capacity for climbing fibers to instruct gain-increase adaptation during contraversive head turns, we used a dual-color optogenetic approach to simultaneously activate climbing fibers and suppress MLIs in *Kit::Cre* mice. Floccular MLIs were transduced with the blue-light sensitive inhibitory opsin *Gt*ACR2 using Cre-dependent AAV, while climbing fibers were transduced with the red-light sensitive excitatory opsin bReaChES by injecting AAV under control of the αCaMKII promoter into the inferior olive (Mathews et al., 2012; Govorunova et al., 2015; Rajasethupathy et al., 2015; Gaffield et al., 2018) (**Figure 4A**). During training sessions, red light pulses were delivered to flocculus to activate climbing fibers, timed to the peak velocity phase of contraversive head turns, while blue light pulses were delivered to suppress MLI activity. This pairing procedure (**Figure 4B,C**) occurred in darkness to avoid visually induced modulation of PC complex spiking. Control training sessions consisting of sinusoidal head motion alone was also obtained in the same mice to account for the effect of darkness-induced habituation on VOR performance (Stahl, 2004).

**Figure 4:**
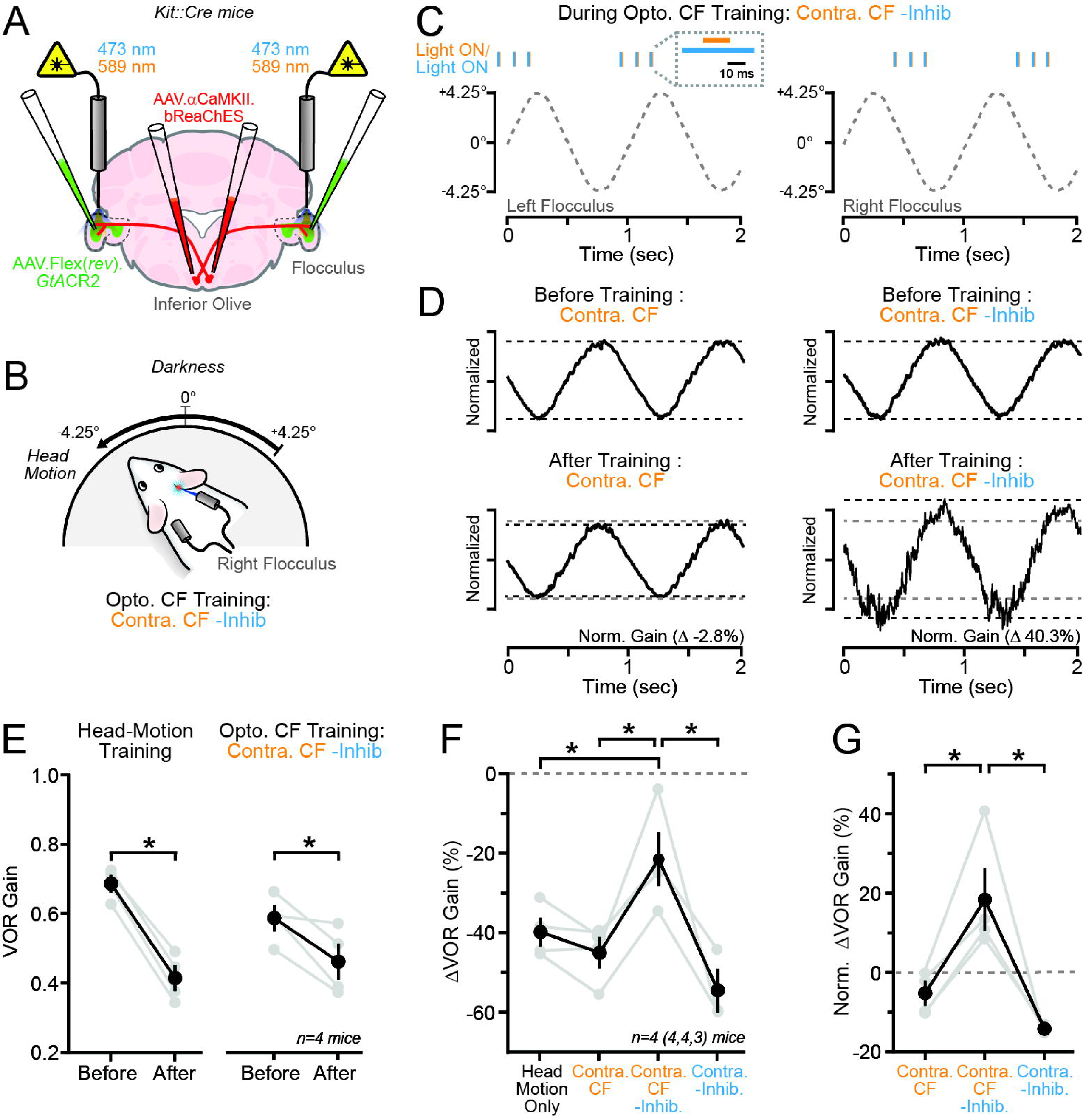
MLI-mediated modulation of optogenetically induced climbing fiber learning. A. In *Kit::Cre* mice, AAV containing bReaChES was injected into the inferior olive to transduce climbing fibers; floccular MLIs were transduced with AAV containing Cre-dependent *Gt*ACR2. Optical fibers delivered light to the each flocculus. B,C. For the training procedure (30 min), climbing fibers were optogenetically activated during the peak velocity phase of contraversive head turns, elicited in darkness, using red light. In some sessions, MLIs were simultaneously suppressed using concurrent pulses of blue light as depicted in panel B. Timing of light pulses (λ = 473 nm and 589 nm; 10 and 8 mW; 25 and 15 ms, respectively) during head turns, with respect to the left and right flocculus, is shown in panel C. D. Average VOR-evoked eye movements measured in the same mouse across different sessions. Left panel: before and after training where climbing fibers were activated during contraversive head turns. Right panel: before and after training that involved climbing fiber activation and MLI activity suppression. Traces in each condition were normalized to that of time-matched, control responses recorded during a session of head turns alone to account for the habituating effect of darkness. E. Summary data of the change in VOR gain for control sessions consisting solely of head turns or when climbing fibers were activated coinciding with MLI activity suppression for the same set of mice (left and right plots: p = *<0.0001 and *0.0215, respectively; Mixed effect model with Bonferroni’s post-tests). F. The change in VOR gain, relative to the within-session baseline response, for the same mice across different training sessions (Head Motion Only vs Contra. CF, p = 0.7881; Head Motion Only vs Contra. CF -Inhib., p = *0.0443; Head Motion Only vs Contra. -Inhib., p = 0.1601; Contra. CF vs Contra. CF -Inhib., p = *0.0124; Contra. CF vs Contra. -Inhib., p = 0.4703; and Contra. CF -Inhib. vs Contra. -Inhib., p = *0.0029; Mixed effects model with Tukey’s post-tests). Note that one mouse was not tested in the Conta. -Inhib. condition. G. Data from panel F, but after normalizing for darkness-induced habituation which was isolated for each mouse in a control session which consisted solely of head turns (Contra CF vs Contra CF -Inhib. *p = 0.0302; Contra CF vs Contra -Inhib., p = 0.4479; and Contra CF -Inhib. vs Contra -Inhib., *p = 0.01188; Mixed effects model with Tukey’s post-tests). In plots, individual mice are denoted in gray with the mean ± SEM; asterisk indicate significance.

As previously reported (Nguyen-Vu et al., 2013; Kimpo et al., 2014; Rowan et al., 2018), the pairing of optogenetic climbing fiber activation with contraversive head turns did not induce an adaptive change to the VOR gain when compared to control sessions of head turns alone (**Figure 4D-G**). However, when we used dual color optogenetics to activate climbing fibers coincident with the suppression of MLIs, pairing with contraversive head turns during training induced a relative increase in the VOR gain when compared to control training sessions (**Figure 4D-G**). The direction and magnitude of the learned response elicited by optogenetic climbing fiber activation in the disinhibited condition resembled that induced by pairing optogenetic climbing fiber activity with ipsiversive head turns (Nguyen-Vu et al., 2013; Kimpo et al., 2014; Rowan et al., 2018) as well as the gain-increase adaptation induced by opposite-direction visual-head motion (see **Supplemental Figure 1B**). Training sessions that included blue light pulses timed to the peak velocity phase of contraversive head turns did not produce a change in VOR gain that was different from control training sessions (**Figure 4F,G**) indicating that the induced learned change was not due to MLI activity suppression alone. Together, these results demonstrate that disinhibiting PCs through MLI activity suppression produces a state change in the floccular circuit that allows for the acquisition of climbing-fiber-instructed learning.

### Learning deficits in the absence of MLI-mediated disinhibition of PCs

We hypothesized that if MLI-mediated inhibition of PCs opposes climbing-fiber-mediated instructive signaling to avoid maladapting movements in mistake-free contexts, then the output of MLIs onto PCs must be relieved during motor errors to facilitate corrective learning. MLIs receive GABA-mediated synaptic input from other MLIs, PCs, and PC-layer interneurons. The inhibition of MLIs from these inputs results in PC disinhibition (Rieubland et al., 2014; Blot et al., 2016; Arlt and Häusser, 2020). To test whether gated cerebellar learning depends on this disinhibitory circuit, we injected AAV containing Cre-dependent, gephyrin-targeted E3 ligase (GFE3; Gross et al., 2016) into the flocculi of adult *Kit::Cre* mice to ablate GABA synapses onto MLIs (**Figure 5A**). In whole-cell recordings obtained from acute slices, GFE3-expressing MLIs showed highly diminished rates of spontaneous inhibitory post-synaptic currents (sIPSCs) relative to MLIs transduced with control constructs (**Figure 5B,C**). This result indicates the functional loss of inhibitory signaling onto MLIs due to GFE3-dependent GABA_A_ receptor de-clustering at inhibitory synapses. However, spontaneous IPSC rates in PCs recorded in areas with a high density of GFE3-expressing MLIs were not different from the rates observed in control-injected animals (**Figure 5D**). Thus, MLI-mediated inhibition of PCs was spared. As GFE3-expressing MLIs cannot be inhibited but continue to inhibit PCs, we conclude that a prominent PC disinhibitory circuit is functionally lost in these animals.

**Figure 5.**
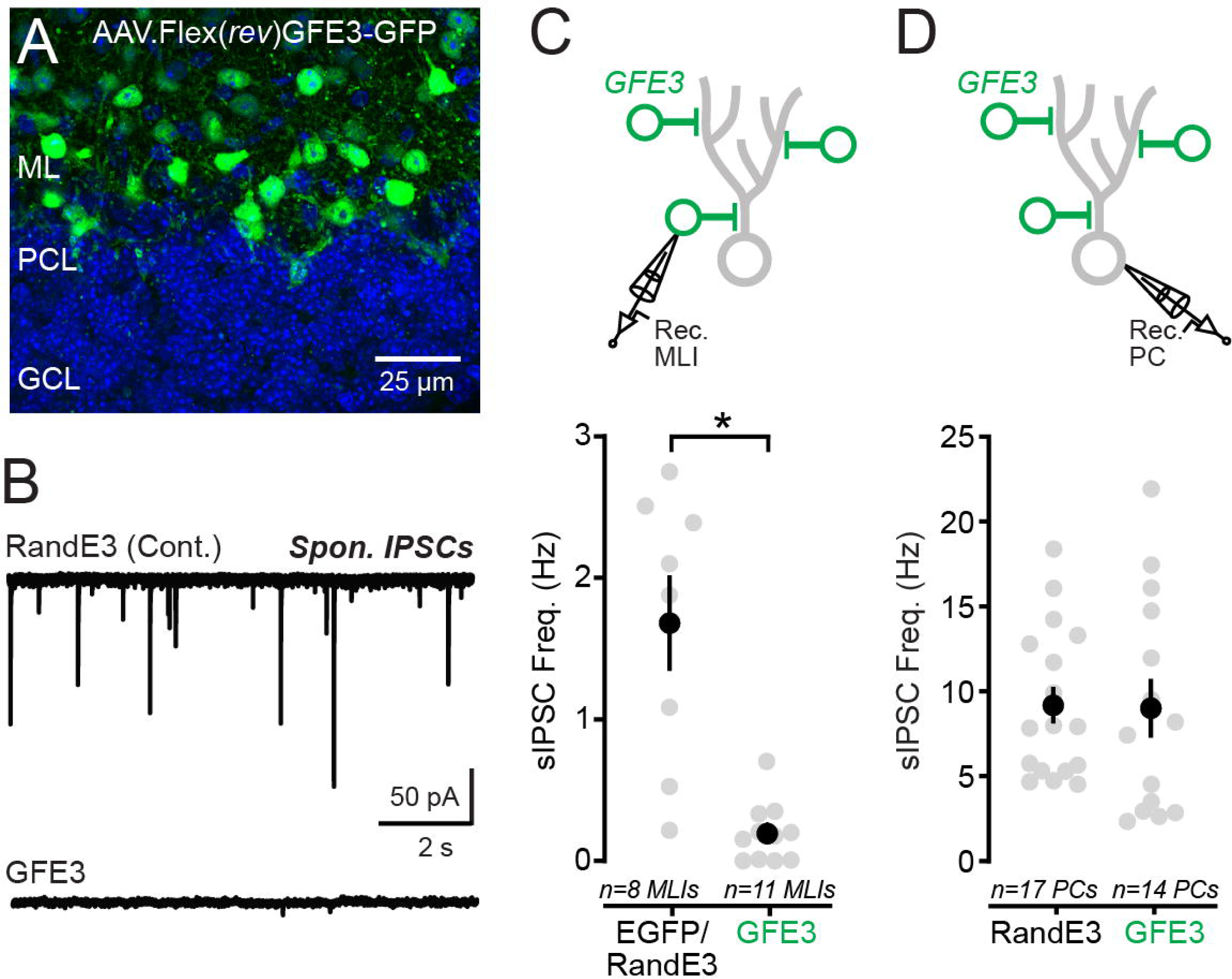
GFE3-mediated ablation of inhibitory synapses onto MLIs. A. Fluorescence image from the cerebellum of a *Kit::Cre* mouse injected with AAV containing Cre-dependent GFE3-eGFP. B. Spontaneous (s)IPSC activity traces from MLIs expressing either a control construct (top) or GFE3 (bottom). C. Comparison of sIPSC rates in floccular MLIs transduced with Cre-dependent AAVs containing control constructs (GFP or RandE3) or GFE3 (p = *<0.0001; unpaired t-test). Circles are measurements from individual cells. D. Summary data showing that sIPSC rates in floccular PCs remain unaffected by the ablation of inhibitory synapses onto MLIs (p = 0.9284; unpaired t-test).

We next assessed the ability of GFE3-expressing mice to adapt their VOR in response to retinal slip errors by pairing sinusoidal head turns with opposite-direction visual motion (**Figure 6A,B**). In contrast to control-injected animals that exhibited gain-increase learning, GFE3-expressing mice failed to adapt the gain of their VOR (**Figure 6C-E**). Importantly, baseline measurements prior to the training procedure showed that VOR-evoked eye movements in GFE3-expressing mice were no different than control-injected animals (1 Hz VOR Gain = 0.82 ± 0.02 and 0.81 ± 0.04, RandE3 and GFE3, respectively p = 0.8989; paired t-test). Thus, ablating inhibitory synapses onto MLIs did not compromise VOR performance in naïve animals, which could lead to learning deficits. These results indicate that, in the absence of a circuit to suppress MLI activity, the mice did not learn to correct their VOR despite receiving retinal slip feedback reporting an under-performance error. However, the same animals learned to adapt their VOR when trained with gain-decrease stimuli (same-direction visual-head motion mismatches) in separate sessions (**Figure 6F,G**). The magnitude of gain-decrease adaptation in GFE3-expressing animals was no different than that acquired in mice injected with control AAV (**Figure 6G,H**), indicating that this form of cerebellar motor learning does not require inhibitory signaling onto MLIs.

**Figure 6.**
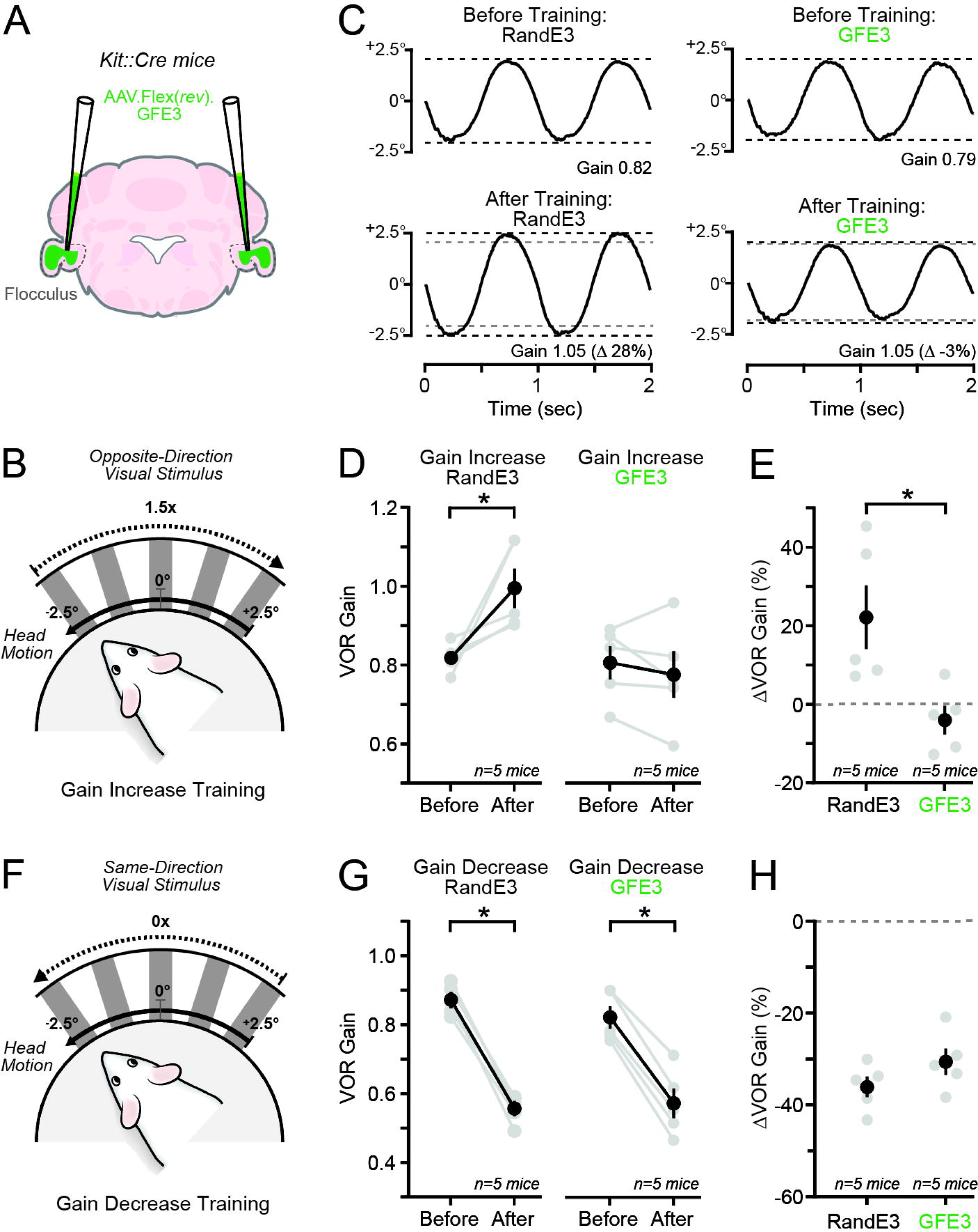
Assessing VOR learning in GFE3-expressing mice. A. Bilateral injections of Cre-dependent AAV into the flocculi of *Kit::Cre* mice were used to transduce MLIs with GFE3. B. Depiction of the gain-increase training stimulus (60 min). C. Average VOR-evoked eye movements recorded in separate mice expressing either a control construct (left) or GFE3 (right), before and after training. D. Alterations in VOR gain observed in groups of control and GFE3-expressing mice for the gain-increase training procedure (left and right plots: p = *0.0141 and >0.99, respectively; ANOVA with Bonferroni’s post-tests). E. Data from panel D presented as the change in VOR gain relative to the baseline response measured at the start of each session for each mouse (p = *0.0188; unpaired t-test). F. Mice were also trained in sessions consisting of the gain-decrease stimulus (60 min). G, H. Similar to panels D and E, except for training with the gain-decrease stimulus (panel G, left and right plots: p = *<0.0001 and *<0.0001, respectively; ANOVA with Bonferroni’s post-tests; panel H, p = 0.1732, unpaired t test). Data in plots are from individual mice are in gray with the mean ± SEM; asterisk denotes significance.

### MLI activity suppression rescues learning in the absence of PC disinhibition

Our results show that in the absence of GABAergic signaling onto MLIs, mice lose their ability to acquire an adaptive increase in VOR gain when confronted by opposite-direction retinal slip errors. An interpretation of this result is that MLI-mediated inhibition of PCs cannot be relieved to allow unopposed climbing-fiber-mediated instructive signaling (Callaway et al., 1995; Gaffield et al., 2018; Rowan et al., 2018). If true, we reasoned that gain-increase VOR learning would be restored in GFE3-expressing mice if MLI activity was extrinsically suppressed during the training procedure because this manipulation would broadly mimic the effect of synaptic inhibition on MLI output, which would result in PC disinhibition.

To test this idea, both flocculi of *Kit::Cre* mice were co-injected with AAVs containing Cre-dependent GFE3 as well as eNphr3.0 (**Figure 7A**). In training sessions including gain-increase stimuli (opposite-direction visual-head motion mismatches), co-injected mice did not adapt the gain of their VOR (**Figure 7E-G**) recapitulating the learning deficit observed in mice expressing GFE3 alone. We then retested the same co-injected mice in separate training sessions but included the optogenetic suppression of MLI activity during the pairing procedure to disinhibit PCs (**Figure 7B,C**). When light pulses were timed to the peak velocity phase of ipsiversive head turns (**Figure 7B,D**), the mice exhibited VOR gain-increase learning (**Figure 7E-G**). Therefore, with MLI activity suppressed, the mice recovered their ability to adapt their VOR in response to retinal slip errors. By contrast, when the light pulses were timed to the peak velocity phase of contraversive head turns, gain-increase learning was still absent (**Figure 7F,G**). This suggests that imposition of PC disinhibition only during ipsiversive head turns was effective at recovering gain-increase learning. We conclude that PC disinhibition mediated by GABAergic signaling onto MLIs is required for the acquisition of context-dependent, error-driven motor learning instructed by the activity of climbing fibers.

**Figure 7.**
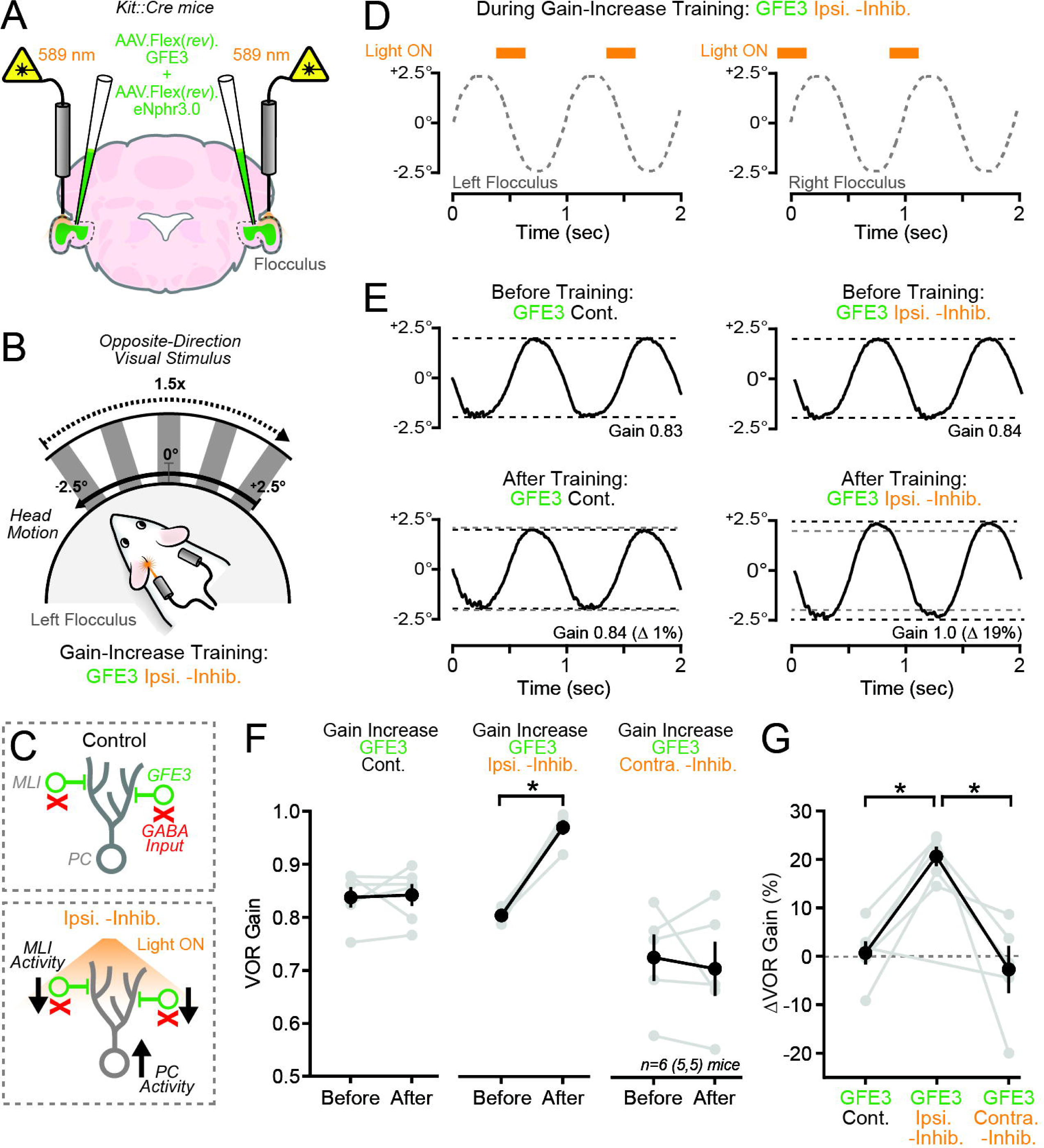
Optogenetic rectification of VOR learning deficits in GFE3-expressing mice. A. Both flocculi of *Kit::Cre* mice were injected with AAVs containing Cre-dependent GFE3 and eNphr3.0. Implanted optical fibers delivered laser light to the transduced cerebellar regions. B. Training involved the gain-increase stimulus while optogenetically suppressing MLI activity (60 min) during ipsiversive (illustrated for the left flocculus in panel B) or contraversive head turns. C. Cartoon depiction of cerebellar circuitry in the control condition with GFE3-expressing MLIs lacking inhibitory input and with optogenetic MLI activity suppression, which disinhibits PCs. Timing of light pulses (l = 589 nm; 8 mW; 240 ms) during the peak velocity phase of ipsiversive head turns, relative to the left and right flocculus, is depicted in panel C. D. Average VOR eye movements from the same GFE3-expressing mouse obtained either in a control session without the optogenetic perturbation (left), or in session incorporating MLI activity suppression during ipsiversive head turns (right). E. Summary data of the change in VOR gain across in the same mice across the different training session conditions (left to right plots: p = 0.9969, *0.0003, and p = 0.8380, Mixed effects model with Bonferroni’s post-tests). Note, two of the mice were only tested in one experimental condition (i.e., either Ipsi. -Inhib. or Contra. -Inhib.). F. Data in panel E presented as the change in VOR gain relative to the baseline response recorded at the start of each session for each mouse (Cont. vs Ipsi. -Inhib., p = *0.0019; Cont. vs Contra. -Inhib., p = 0.7395; and Contra. -Inhib. vs Ipsi. -Inhib., p = *0.0007; Mixed effects model with Tukey’s post-tests). Data in plots are from individual mice are in gray with the mean ± SEM; asterisk denotes significance.

## Discussion

By manipulating the activity of floccular MLIs during the VOR, our current study sheds light on the mechanisms underlying supervised cerebellar learning. In contrast to the view that climbing fiber activity reliably triggers motor adaptation, our results point to a layered process by which inhibition from MLIs antagonizes the potency of climbing fibers to sculpt movement through plasticity induction unless PCs become disinhibited through an MLI-mediated pathway. Such dynamic gating by MLIs helps ensure that climbing-fiber-mediated learning is instantiated based on error relevance, thus avoiding the induction of maladaptive plasticity that alters otherwise well performed behaviors.

Our functional recordings revealed increased complex spiking in floccular PCs when head turns were elicited in association with a non-moving visual stimulus. Despite the modulation of climbing-fiber-evoked activity, the mice did not adapt their VOR gain, likely because the behavior was performed with little retinal slip. Thus, climbing fiber activity was not exclusively predictive of whether the mice would learn to adjust their motor performance. These results are consistent with prior observations showing that climbing fibers fire during well performed movements and that climbing fiber activity is not always correlated with learned performance changes (Kimpo et al., 2014; Heffley et al., 2018; Kostadinov et al., 2019). However, by optogenetically suppressing MLI activity during the same VOR context, we found that the mice learned to increase their VOR gain even though performance errors were mostly absent. This implies that in the disinhibited condition, climbing fibers become supra-potent for instructing changes to the VOR because gain-increase learning is mediated by these olivary inputs (Kimpo et al., 2014). We substantiated this idea using dual-color optogenetics, causally demonstrating that MLI activity suppression unlocks the capacity of climbing fibers to induce gain-increase learning during a context in which climbing fibers are normally ineffective at instructing a VOR performance change (Nguyen-Vu et al., 2013; Rowan et al., 2018). We conclude that MLIs gate climbing-fiber-mediated instructive signaling during correctly performed movements.

We propose that in the absence of inhibitory gating of climbing fiber input by MLIs, PCs would continuously integrate instructive signals during well performed movements, resulting in maladaptive plasticity induction. Unregulated climbing-fiber-induced plasticity is known to be pathological (Piochon et al., 2014; Nguyen-Vu et al., 2017), therefore, plasticity constraint is likely necessary for the healthy cerebellum. We suggest that an MLI-mediated gating mechanism serves this function. In many brain regions, regulatory pathways for plasticity constraint commonly involve inhibitory interneurons whose GABAergic output dampens the responsiveness of their postsynaptic principal cell targets to excitatory input (Paulsen and Moser, 1998; Larkum et al., 1999; Tang et al., 2021). The cerebellum appears to employ similar neural circuit mechanisms to gate plasticity induction and maladaptive motor learning.

Our findings support the concept that MLI gating of climbing-fiber-mediated instructive signaling not only opposes learned behavioral adjustments in the absence of motor errors but also ensures that movements are appropriately adapted when mistakes occur. The murine vestibulo-cerebellum is optimized to enhance VOR eye movements during ipsiversive head turns (Voges et al., 2017). Therefore, plasticity constraint during contraversive head turns, but not ipsiversive head turns, is likely required to prevent isometric strengthening of both eye movement directions, which could result in no net gain change. We observed the absence of a VOR gain change, despite that mice were experiencing opposite-direction retinal slip, when MLI activity was suppressed during contraversive head turns, a result consistent with such an isometric effect. Similarly, MLI gating of climbing-fiber-mediated instructive signaling during same-direction retinal slip may prevent gain-increase learning that would counteract the necessary adaptive weakening of VOR gain to correct an erroneously over-performing VOR. Our results support this conclusion, as we found that optogenetic MLI activity suppression weakened VOR gain-decrease learning when timed to ipsiversive head turns.

Climbing fibers have long been implicated as a source of instructive signals in the cerebellar cortex that adapt erroneously performed movements (De Zeeuw et al., 1998; Ito, 2013). In addition to evoking somatic complex spikes in PCs, climbing fibers also elicit simultaneous bursts of dendritic calcium action potentials (Llinas and Sugimori, 1980; Kitazawa et al., 1998; Davie et al., 2008; Ozden et al., 2009). This PC calcium response is of particular interest in the instructive signaling pathway because intracellular calcium elevation induces plasticity at parallel-fiber-to PC synapses (Finch et al., 2012; Gaffield et al., 2019; Bonnan et al., 2021; Fanning et al., 2021; Silva et al., 2022). We posit that during well performed movements, MLIs exert control over climbing-fiber-mediated instructive signaling by inhibiting the PC dendritic calcium response to climbing fiber input and that this regulation is relieved by a disinhibitory mechanism during motor errors. Prior investigations have shown that MLI activation reduces climbing-fiber-evoked dendritic calcium signaling in PCs, impeding LTD induction at co-active parallel fiber synapses, and preventing VOR gain increase learning to retinal slip errors (Ekerot and Kano, 1985; Callaway et al., 1995; Rowan et al., 2018). Furthermore, blocking MLI activity increases climbing-fiber-evoked PC calcium responses during well performed behaviors (Gaffield et al., 2018). Future work examining dendritic calcium responses in individual floccular PCs, rather than their population (Rowan et al., 2018; Fanning et al., 2021), will help delineate if dendritic calcium signal magnitude varies dependent on error context during VOR performance, and whether MLI-mediated inhibition plays a determinate role in this process.

We discovered that ablating GABAergic synapses onto MLIs prevented gain-increase VOR learning. This suggests that inhibitory activity within the MLI ensemble and the consequent disinhibition of PCs is a crucial permissive step in error-driven instructive signaling by climbing fibers. To support this mechanism, we propose that the response properties of MLIs, and the net overall effect of their activity on PC excitation, varies with the context of motor performance. The MLI population, which is composed of molecularly and functionally distinct subtypes (Kozareva et al., 2021), is highly responsive to movements, exhibiting both increases and decreases in spiking within their ensemble during motor output (Jelitai et al., 2016; Astorga et al., 2017; Chen et al., 2017; Gaffield and Christie, 2017). However, whether MLIs are differentially engaged based on error context, and the net effect of their activity on PC excitability is still unknown. Interestingly, reciprocal synaptic connectivity between any two MLIs is rare (Kondo and Marty, 1998; Christie et al., 2011; Rieubland et al., 2014) and MLIs appear to favor a structured, unidirectional inhibitory connectivity in their network. An interesting possibility is that one MLI subtype may preferentially inhibit a different, PC-targeting MLI subtype(s), and that this disinhibitory circuit is specifically engaged during motor errors. Additionally, PC disinhibitory circuits, routed through inhibitory connections onto MLIs may also involve PC-layer interneurons and PCs, that latter through feedback inhibition (Halverson et al., 2022; Osorno et al., 2022).

In summary, the results of our current study challenge a well-established theory of cerebellar function, which posits that the coincident activity of parallel fibers and climbing fibers alone is necessary for motor learning (Marr, 1969; Albus, 1971; Ito, 1986). Instead, we propose that associative learning occurs when these inputs fire conjunctively in the absence of inhibitory drive from PC-targeting MLIs. Our findings indicate that this synergistic regulation can prevent maladapting well-performed behaviors and biases learning to that based on error relevance. Our study thus presents a paradigm shift in the understanding of cerebellar circuit function in motor learning behavior.

## Acknowledgements

The authors thank Samantha Amat and Audrey Bonnan (Christie Lab) for their helpful input and assistance with the project. This work was supported by National Institutes of Health Grants NS115610 (D.B.A) and NS118401, NS105958, and NS123933 (J.M.C).

## Author Contributions

Conceptualization, J.M.C.; Investigation, K.Z. and Z.Y.; Analysis, K.Z., Z.Y., and M.A.G.; Resources, G.G.G. and D.B.A.; Writing, J.M.C.; Funding Acquisition, J.M.C.; Supervision, J.M.C.

## Figure Legends

**Supplemental Figure 1. Modulation of VOR learning through gain-increase and gain-decrease training**

A,B. Left panel: Gain-increase training involved pairing of sinusoidal head turns with opposite-direction visual motion (60 min). Right panel: Average VOR-evoked eye movements from a mouse before and after training. The gray dotted line in the bottom eye trace indicates the gain measurement obtained immediately prior to training.

C. The effect of the training procedure on the VOR gain across mice (p = *0.0117; paired t-test).

D. Summary of the effect of. Data are from individual mice along with the mean ± SEM.

**Supplemental Figure 2. Effect of MLI activity suppression on VOR-evoked PC population responses**

A. AAVs containing Cre-dependent eNphr3.0, FlpO under control of the PC L7 promoter, and FlpO-dependent GCaMP6f were injected into the left flocculi of Kit::Cre mice. Implanted optical fibers targeted the infected region of cerebellum.

B. A histological image from an injected mouse shows GCaMP6f and eNphr3.0 expression in PCs and MLIs, respectively.

C. Sinusoidal head turns were paired with a stationary visual surround (i.e., the gain-stable training stimulus) while measuring Ca^2+^ activity from the PC population using photometry in conjunction with optogenetics.

D. In an example mouse, trial averaged Ca^2+^ activity in the PC population in the control condition and during MLI activity suppression during ipsiversive head turns. The artifact from the laser pulse was removed for clarity.

E. Summary showing the effect of MLI activity suppression on the PC population Ca^2+^ response activity during optogenetic MLI suppression compared to the control response without the optogenetic stimulus. The data are from individual mice in gray along with the mean ± SEM.

## Methods

### Animals

In our study, we used male and female heterozygous *Kit::Cre* mice as well as C57Bl/6J animals. The mean age of animals for *ex vivo* slice recording was 8 weeks (range = 6-15 weeks); the mean age for *in vivo* behavior experiments was 15 weeks (range = 8-23 weeks). All experimental procedures were approved by the Max Planck Florida Institute for Neuroscience Animal Care and Use Committee (IACUC).

### Surgical procedures

To perform stereotactic surgeries, mice were immobilized using ear bars and held head-fixed on a platform (David Kopf Instruments). The mice were continuously anesthetized with isoflurane (1.8-4.0%) and their body temperature was maintained at 37°C using a heating plate with biofeedback from a rectal thermometer. Ophthalmic ointment was applied to protect the eyes. Local anesthesia was induced with a subcutaneous injection of a lidocaine/bupivacaine cocktail into the scalp.

For viral injections targeting the flocculi, a small incision was made in the scalp and a craniotomy (<0.5 mm in diameter) was created above the cerebellum to allow access for a microinjection pipette. The microinjection coordinates (in mm from Bregma) were M/L=±2.33; A/P=-5.68; D/V = 2.7-2.8 mm (angle of 5°). For targeting the inferior olive, the micropipette was inserted through a surgical opening between the foramen magnum and C1 vertebra (angle of 62°). Viral solutions were injected at a slow rate (∼0.05 - 0.1 μL/min) by pressure, for a total volume of 0.2-0.5 μL per site. The microinjection pipette was held in place (8-10 min) before withdrawal to ensure proper diffusion of the viral vector. The incision was closed using sutures. Animals received analgesia through injection of carprofen and buprenorphine SR LAB (5 mg/kg and 0.35 mg/kg doses, respectively). Custom-engineered head-posts, fabricated from stainless steel, were affixed to the skull of viral-injected mice used for behavior experiments. In some of these animals, optical fibers (200 µm, NA 0.22 with Ø 1.25 mm ferrules; Thorlabs) were then bilaterally implanted in each side of the cerebellum to deliver light to the virally infected region of flocculus with the following coordinates: M/L = ± 3.35 mm; A/P = 5.65 mm; D/V = 1.9 ± 0.1 mm (angle of −16°). A unilateral implant was used for photometry (MFC_400/430-0.48_MF1.25_FLT, Doric Lenses Inc.) targeting a similar location (M/L = 2.35 mm; ; A/P = 5.65 mm; D/V = 3 ± 0.1 mm; angle of −10°). The head posts and optical fibers were secured to the exposed skull next using dental cement (Metabond; Parkell).

AAVs used to transduce MLIs included AAV5-EF1α-Flex(*loxP*)rev-eNphr3.0-EYFP, AAV9-EF1α-Flex(*loxP*)rev-GFE3-eGFP, AAV1-EF1α-Flex(*loxP*)rev-RandE3-2A-GFP, AAV1-EF1α-Flex(*loxP*)rev-EGFP, AAV1-EF1α-Flex(*loxP*)rev-*Gt*ACR2, and AAV5-EF1α-Flex(*loxP*)rev-eNphr3.0-HA. AAVs used to transduce PCs included AAV1-Pcp2.6-FlpO and AAv1-CAG-Flex(*FRT*)rev-GCaMP6f (Nitta et al., 2017; Gaffield et al., 2019). For transducing inferior olive neurons, we used AAV5-CaMKIIα-bReaChES-EYFP. All AAVs were obtained from vector core facilities of either the University of Pennsylvania or University of North Carolina, or custom packaged at ViGene (Rockville, MD) and were injected at a titer of ≥10^12^ vg/mL.

### VOR testing

To measure the VOR, mice were head-restrained using surgically implanted head-posts on a custom-made, fully enclosed apparatus for controlled light exposure. The setup included a motorized rotation stage (T-RSW60C, Zaber Technology) for passive head turns, and a camera with a machine-vision lens (MVL6X12Z; Thorlabs) aimed at the left eye to capture VOR-evoked movements. The camera was attached to the rotation stage perimeter on a rail, allowing its position to be equidistant from the eye’s center. A commercial video system (ETL-200; ISCAN) was used to compute relative eye movements by tracking the pupil. The system provided horizontal pupil position and diameter as linearized voltage signals (120 Hz). Three infrared (IR) LEDs illuminated the eye, with two affixed to the rotation stage below the eye and a third (M940L3; Thorlabs) attached to the top of the camera which provided the reference corneal reflection (*CR*) during analysis. Two LCD monitors on either side of the apparatus provided visual stimuli, with LED backlights controlled by an external TTL command. DAQ boards (PXIe-6356 and PXIe-6383; National Instruments) facilitated computer interfacing with the rotation stage, eye metrics, and visual stimuli on the monitors. Custom-written LabVIEW routines (National Instruments) managed software control.

VOR performance was assessed in darkness. Before testing, pilocarpine (2% ophthalmic drops; Patterson Veterinary Supply) was applied (< 1 min) to the eye to limit pupil dilation for accurate tracking in darkness. For the visual-vestibular pairing procedure during training, a high-contrast grating of black and white vertical stripes was displayed on the monitors. Gain-increase or gain-decrease training had the grating move out-of-phase or in-phase with the vestibular stimulus, respectively. The visual stimulus was kept stationary during gain-stable training. Darkness-isolated VOR performance was gauged before training and retested immediately afterward. One training condition was tested per session, with one session per day, and at least 48 hours between sessions for all conditions. Training sessions were randomized and baseline VOR performance across sessions showed no significant difference (ANOVA with Bonferroni post-hoc tests; P > 0.05). Mice were familiarized with the VOR apparatus for ∼15 minutes before training, improving eye-tracking quality.

Angular eye position was derived using a previously published method (Stahl et al., 2000) which included a calibration procedure for estimating the distance from the pupil plane to the corneal curvature center (*Rp*). Calibration measurements were made for several different pupil diameters to accommodate for a range of diameters encountered during an experiment. Camera translation on the rail (± 10°) allowed *Rp* calibration by measuring the pupil position (*P*). Motion-induced artifacts were corrected using the corresponding *CR* position. *Rp* was calculated across pupil diameters as: *Rp = Δ/sin (20°)*, and angular eye position (*Ep*) was determined using *Ep = arcsin [(P_1_-CR_1_)-(P_2_-CR_2_)/Rp]*. VOR gain was computed as the amplitude ratio between eye and stage. To facilitate comparisons across animals and conditions, changes in VOR gain were also calculated as the percentage difference in VOR performance after training relative to the baseline measurement before training (ΔVOR).

### In vivo optogenetics and photometry

For photo-suppressing eNphr3.0-expressing MLIs in behaving mice, we used a DPSS laser centered on 589 nm (± 1 nm) (CNI Optoelectronics Tech; MGL-F-589-200mW). For suppressing *Gt*ACR-expressing MLIs or exciting ChR2-expressing climbing fibers, we used a DPSS laser centered on 473 nm (± 1 nm) (CNI Optoelectronics Tech; MBL-F-473-200mW). To deliver light from the lasers to the implanted optical fibers of experimental animals, light from each laser was first passed through a shutter (NS15B1T0-ED; Vincent Associates) and then was split into two separate lines using a wave plate (WPH05M-588 and WPH05M-473, respectively; Thorlabs) and beamsplitter cube (PBS101; Thorlabs). Each line was then sent through an acousto-optic modulator (AOM) (AA Opto-Electronic; MTS110-A3-VIS), which allowed for the independent modulation of laser power at a rapid rate (<1 kHz). From each AOM, separate lines of 589 nm and 473 nm laser light were combined using a fiber compatible dichroic mirror (Mini-cube E[470]_E[590]; Doric Lenses) and was then launched into fiber ports (PAF-X-11-A; Thorlabs); attached patch cables (MFP_200/240/900-0.22; Doric Lenses) delivered the light to the optical fiber implants targeting each flocculi. Laser pulses for optogenetics were triggered by the position of the stage. Thus, light stimuli were timed to head position.

For photometry, blue light from an LED (centered at λ = 461 nm; M470F3; Thorlabs) was launched into a fiber-compatible dichroic mirror mount (Fluorescence Mini-cube FMC6_E1[400-410]_F1[420-450]_E2[460-490]_F2[500-540]_O[570-650]_S; Doric Lenses). A patchcord (MFP_400/430/1100-0.48_FC-SMA, Doric Lenses Inc.) delivered excitation light (∼20-40 µW) to the implanted optical fiber targeting the transduced region of the flocculus. Emitted fluorescence was collected through the same implanted optical fiber and detected using a femtowatt photoreceiver (Model 2151; Newport) and a DAQ board (PXIe-6363; National Instruments). After binning samples, the effective rate of activity measurements was 25 Hz. Red laser light from a DPSS laser (λ = 589 nm) was launched into the same dichroic mirror mount from a fiber port (PAF-X-11-A; Thorlabs) to support simultaneous optogenetic manipulations during photometry recording. Prior to the fiber port, the laser light was gated by a fast shutter (NS15B1T0-ED; Vincent Associates) and modulated by an AOM (AA Opto-Electronic; MTS110-A3-VIS). Custom-written software in LabVIEW (National Instruments) controlled the system. Trials that included visual and optogenetic pairing were presented in a randomized manner (120 s each) for each mouse with a period of separation (120 s) between each condition. Baseline fluorescence for calculating ΔF/F was simply mean fluorescence prior to the optogenetic stimulus.

### In vivo electrophysiology

The flocculus was targeted for electrophysiological recordings through a surgically prepared craniotomy, using the same coordinates as for optical fiber implants. A silver wire, placed at the craniotomy site, served as a ground signal, and the opening was bathed in saline during recording. A silicon probe (A1×32-Poly3-5mm-25s-177, Neuronexus; H6B and ASSY-79 P-1, Cambridge Neurotech) was carefully lowered to the recording site using a manual stereotactic micromanipulator (David Kopf Instruments). The silicon probe was connected to an amplifier (RHD2132, Intan Technologies) and read by a controller interface (RHD200, Intan Technologies) at a sampling rate of 20 kHz. Data acquisition was performed using commercial software (Intan Technologies). Silicon probe data were automatically sorted using the Kilosort algorithm (Pachitariu et al., 2016), followed by manual curation with Phy2 software (https://github.com/cortex-lab/phy). PC units were unambiguously identified by the presence of accompanying complex spikes in the recordings, which were analyzed based on alignment with head position during passive vestibular stimulation.

### Acute slice preparation and electrophysiology

To prepare brain slices, we first anesthetized mice with ketamine/xylazine (intraperitoneal injection; 20mg/ml; 2mg/ml, respectively). Next, we transcardially perfused the mice with a cold (∼4°C) saline solution containing (in mM) 87 NaCl, 25 NaHO_3_, 2.5 KCl, 1.25 NaH_2_PO_4_, 7 MgCl_2_, 0.5 CaCl_2_, 10 glucose, and 75 sucrose to rapidly chill the brain. The cerebellum was immediately isolated by dissection and slices (200 μm) were sectioned using a vibrating microtome (VT1200S, Leica Biosystems) in an icy saline solution of the same composition. The slices were then transferred to an incubation chamber containing (in mM) 128 NaCl, 26.2 NaHO_3_, 2.5 KCl, 1 NaH_2_PO_4_, 1.5 CaCl_2_, 1.5 MgCl_2_ and 11 glucose and maintained at 34°C for 40 min. They remained at room temperature thereafter. Slices were continuously perfused with this saline solution during whole-cell recording with the addition of 20 μM NBQX and 20 μM (R)-CPP (Tocris) to isolate inhibitory synaptic activity; TTX (1 μM) was sometimes added to isolate miniature synaptic events. All solutions were equilibrated and maintained with carbogen gas (95% O_2_/5% CO_2_).

To target neurons for whole-cell recording, we used visual guidance from gradient-contrast video-microscopy. PCs were identified by their characteristic morphology, and MLIs were identified by their location in the molecular layer. Virally transduced MLIs were identified by marker protein fluorescence. We did not discriminate MLIs subtypes in our recordings. For both PCs and MLIs, we used borosilicate patch pipettes containing an intracellular solution composed of (in mM) 150 CsCl, 10 HEPES, 1 EGTA, 0.1 CaCl_2_. 4.6 MgCl_2_, 2 MgATP, and 0.3 NaGTP (pH = 7.3). The open tip resistance of the patch pipettes was 2-4 MΩ for PCs and 3-5 MΩ for MLIs. The membrane potential of both cell types was maintained at a hyperpolarized level (∼ –80 mV) to better detect inhibitory synaptic events. Electrophysiological measurements were obtained using Multiclamp 700B amplifiers (Molecular Devices) with the analog signals filtered at 2-10 kHz and sampled at 20-50 kHz using a Digidata 1440 digitizer (Molecular Devices). Data was collected using pClamp 10 software (Molecular Devices). Pipette capacitance was neutralized in all recordings, and electrode series resistance compensated using bridge balance in current-clamp mode. Liquid junctional potentials, determined to be −20 mV, were corrected in our recordings.

### Histology

For post-hoc examination of virally-transduced brain tissue, mice were anesthetized using a ketamine and xylazine cocktail (100 mg/kg and 10 mg/kg, respectively). The mice were then transcardially perfused with 0.1 M phosphate buffer (PB), followed by 4% paraformaldehyde (PFA) in PB. The cerebellum was dissected and post-fixed in the same PFA solution for approximately 12 hours at 4°C. After washing the fixed tissue several times with PB, it was embedded in 4% agar and sectioned into 100-µm slices using a vibraslicer (VT1200S, Leica Biosystems). The slices were mounted on glass slides with anti-fade media (#S36963, Thermofisher Scientific).

For visualizing expression of AAV5-EF1a-Flex(*loxP*)-eNphr3.0-HA injected mice, the sectioned slices were incubated for one hour at room temperature in blocking solution (10% normal goat serum and 0.3% Titron X-100 in 1x TBS) and incubated overnight in polyclonal rabbit anti-HA (1:1,000, #ab9110, Abcam) at 4°C. The slices were then washed in 1x TBS and incubated for 1 hour at room temperature in AlexaFluor 633 goat anti-rabbit (1:1,000 #A-21070, Thermofisher Scientific) and mounted on slides after repeated PBS washes.

Fluorescence images were captured using a confocal scanning microscope (Zeiss LSM880). Maximum-intensity projection images were generated from stacks of scanned tissue (Z-steps of approximately 30 µm) using ZEN Light software (Zeiss).

### Statistics

Population data are presented as mean ± SEM and include the number of samples in either the figure or figure legend. Differences were considered significant at α values of P < 0.05 using paired or unpaired t-tests. When suitable, group data were compared using ANOVA or Mix effects model, and significance between groups was determined using either Tukey’s and Bonferroni’s post-hoc comparison tests as appropriate.

## References

Albus JS (1971) A theory of cerebellar function. Math Biosci 10:25–61.

Amat SB, Rowan MJM, Gaffield MA, Bonnan A, Kikuchi C, Taniguchi H, Christie JM (2017) Using c-kit to genetically target cerebellar molecular layer interneurons in adult mice. PloS one 12:e0179347.

Arlt C, Häusser M (2020) Microcircuit Rules Governing Impact of Single Interneurons on Purkinje Cell Output In Vivo. Cell reports 30:3020–3035.e3023.

Astorga G, Li D, Therreau L, Kassa M, Marty A, Llano I (2017) Concerted Interneuron Activity in the Cerebellar Molecular Layer During Rhythmic Oromotor Behaviors. The Journal of neuroscience : the official journal of the Society for Neuroscience 37:11455–11468.

Blot A, de Solages C, Ostojic S, Szapiro G, Hakim V, Lena C (2016) Time-invariant feed-forward inhibition of Purkinje cells in the cerebellar cortex in vivo. The Journal of physiology 594:2729–2749.

Bonnan A, Rowan MMJ, Baker CA, Bolton MM, Christie JM (2021) Autonomous Purkinje cell activation instructs bidirectional motor learning through evoked dendritic calcium signaling. Nature communications 12:2153.

Boyden ES, Katoh A, Raymond JL (2004) Cerebellum-dependent learning: the role of multiple plasticity mechanisms. Annual review of neuroscience 27:581–609.

Boyden ES, Katoh A, Pyle JL, Chatila TA, Tsien RW, Raymond JL (2006) Selective engagement of plasticity mechanisms for motor memory storage. Neuron 51:823–834.

Brunel N, Hakim V, Isope P, Nadal JP, Barbour B (2004) Optimal information storage and the distribution of synaptic weights: perceptron versus Purkinje cell. Neuron 43:745–757.

Callaway JC, Lasser-Ross N, Ross WN (1995) IPSPs strongly inhibit climbing fiber-activated [Ca2+]i increases in the dendrites of cerebellar Purkinje neurons. The Journal of neuroscience : the official journal of the Society for Neuroscience 15:2777–2787.

Chen S, Augustine GJ, Chadderton P (2017) Serial processing of kinematic signals by cerebellar circuitry during voluntary whisking. Nature communications 8:232.

Christie JM, Chiu DN, Jahr CE (2011) Ca(2+)-dependent enhancement of release by subthreshold somatic depolarization. Nature neuroscience 14:62–68.

Davie JT, Clark BA, Hausser M (2008) The origin of the complex spike in cerebellar Purkinje cells. The Journal of neuroscience : the official journal of the Society for Neuroscience 28:7599–7609.

De Zeeuw CI, Simpson JI, Hoogenraad CC, Galjart N, Koekkoek SK, Ruigrok TJ (1998) Microcircuitry and function of the inferior olive. Trends in neurosciences 21:391–400.

Dizon MJ, Khodakhah K (2011) The role of interneurons in shaping Purkinje cell responses in the cerebellar cortex. The Journal of neuroscience : the official journal of the Society for Neuroscience 31:10463–10473.

Ekerot CF, Kano M (1985) Long-term depression of parallel fibre synapses following stimulation of climbing fibres. Brain research 342:357–360.

Fanning AS, Shakhawat AM, Raymond JL (2021) Population calcium responses of Purkinje cells in the oculomotor cerebellum driven by nonvisual input. Journal of neurophysiology 126:1391–1402.

Finch EA, Tanaka K, Augustine GJ (2012) Calcium as a trigger for cerebellar long-term synaptic depression. Cerebellum (London, England) 11:706–717.

Gaffield MA, Christie JM (2017) Movement Rate Is Encoded and Influenced by Widespread, Coherent Activity of Cerebellar Molecular Layer Interneurons. The Journal of neuroscience : the official journal of the Society for Neuroscience 37:4751–4765.

Gaffield MA, Bonnan A, Christie JM (2019) Conversion of Graded Presynaptic Climbing Fiber Activity into Graded Postsynaptic Ca(2+) Signals by Purkinje Cell Dendrites. Neuron 102:762–769.e764.

Gaffield MA, Rowan MJM, Amat SB, Hirai H, Christie JM (2018) Inhibition gates supralinear Ca(2+) signaling in Purkinje cell dendrites during practiced movements. eLife 7.

Ghelarducci B, Ito M, Yagi N (1975) Impulse discharges from flocculus Purkinje cells of alert rabbits during visual stimulation combined with horizontal head rotation. Brain research 87:66–72.

Govorunova EG, Sineshchekov OA, Janz R, Liu X, Spudich JL (2015) NEUROSCIENCE. Natural light-gated anion channels: A family of microbial rhodopsins for advanced optogenetics. Science (New York, NY) 349:647–650.

Graf W, Simpson JI, Leonard CS (1988) Spatial organization of visual messages of the rabbit’s cerebellar flocculus. II. Complex and simple spike responses of Purkinje cells. Journal of neurophysiology 60:2091–2121.

Gross GG, Straub C, Perez-Sanchez J, Dempsey WP, Junge JA, Roberts RW, Trinh le A, Fraser SE, De Koninck Y, De Koninck P, Sabatini BL, Arnold DB (2016) An E3-ligase-based method for ablating inhibitory synapses. Nature methods 13:673–678.

Halverson HE, Kim J, Khilkevich A, Mauk MD, Augustine GJ (2022) Feedback inhibition underlies new computational functions of cerebellar interneurons. eLife 11.

Heffley W, Song EY, Xu Z, Taylor BN, Hughes MA, McKinney A, Joshua M, Hull C (2018) Coordinated cerebellar climbing fiber activity signals learned sensorimotor predictions. Nature neuroscience 21:1431–1441.

Ito M (1982) Cerebellar control of the vestibulo-ocular reflex--around the flocculus hypothesis. Annual review of neuroscience 5:275–296.

Ito M (1986) Long-term depression as a memory process in the cerebellum. Neuroscience research 3:531–539.

Ito M (2013) Error detection and representation in the olivo-cerebellar system. Frontiers in neural circuits 7:1.

Jelitai M, Puggioni P, Ishikawa T, Rinaldi A, Duguid I (2016) Dendritic excitation-inhibition balance shapes cerebellar output during motor behaviour. Nature communications 7:13722.

Kepecs A, Fishell G (2014) Interneuron cell types are fit to function. Nature 505:318–326.

Kimpo RR, Rinaldi JM, Kim CK, Payne HL, Raymond JL (2014) Gating of neural error signals during motor learning. eLife 3:e02076.

Kitamura K, Hausser M (2011) Dendritic calcium signaling triggered by spontaneous and sensory-evoked climbing fiber input to cerebellar Purkinje cells in vivo. The Journal of neuroscience : the official journal of the Society for Neuroscience 31:10847–10858.

Kitazawa S, Kimura T, Yin PB (1998) Cerebellar complex spikes encode both destinations and errors in arm movements. Nature 392:494–497.

Knudsen EI (1994) Supervised learning in the brain. The Journal of neuroscience : the official journal of the Society for Neuroscience 14:3985–3997.

Kondo S, Marty A (1998) Synaptic currents at individual connections among stellate cells in rat cerebellar slices. The Journal of physiology 509 (Pt 1):221–232.

Kostadinov D, Beau M, Pozo MB, Hausser M (2019) Predictive and reactive reward signals conveyed by climbing fiber inputs to cerebellar Purkinje cells. Nature neuroscience.

Kozareva V, Martin C, Osorno T, Rudolph S, Guo C, Vanderburg C, Nadaf N, Regev A, Regehr WG, Macosko E (2021) A transcriptomic atlas of mouse cerebellar cortex comprehensively defines cell types. Nature 598:214–219.

Larkum ME, Zhu JJ, Sakmann B (1999) A new cellular mechanism for coupling inputs arriving at different cortical layers. Nature 398:338–341.

Letzkus JJ, Wolff SB, Luthi A (2015) Disinhibition, a Circuit Mechanism for Associative Learning and Memory. Neuron 88:264–276.

Llinas R, Sugimori M (1980) Electrophysiological properties of in vitro Purkinje cell dendrites in mammalian cerebellar slices. The Journal of physiology 305:197–213.

Marr D (1969) A theory of cerebellar cortex. The Journal of physiology 202:437–470.

Mathews PJ, Lee KH, Peng Z, Houser CR, Otis TS (2012) Effects of climbing fiber driven inhibition on Purkinje neuron spiking. The Journal of neuroscience : the official journal of the Society for Neuroscience 32:17988–17997.

Mittmann W, Koch U, Hausser M (2005) Feed-forward inhibition shapes the spike output of cerebellar Purkinje cells. The Journal of physiology 563:369–378.

Nguyen-Vu TB, Zhao GQ, Lahiri S, Kimpo RR, Lee H, Ganguli S, Shatz CJ, Raymond JL (2017) A saturation hypothesis to explain both enhanced and impaired learning with enhanced plasticity. eLife 6.

Nguyen-Vu TD, Kimpo RR, Rinaldi JM, Kohli A, Zeng H, Deisseroth K, Raymond JL (2013) Cerebellar Purkinje cell activity drives motor learning. Nature neuroscience 16:1734–1736.

Nitta K, Matsuzaki Y, Konno A, Hirai H (2017) Minimal Purkinje Cell-Specific PCP2/L7 Promoter Virally Available for Rodents and Non-human Primates. Molecular therapy Methods & clinical development 6:159–170.

O’Donoghue DL, King JS, Bishop GA (1989) Physiological and anatomical studies of the interactions between Purkinje cells and basket cells in the cat’s cerebellar cortex: evidence for a unitary relationship. The Journal of neuroscience : the official journal of the Society for Neuroscience 9:2141–2150.

Osorno T, Rudolph S, Nguyen T, Kozareva V, Nadaf N, Macosko EZ, Lee W-CA, Regehr WG (2021) Candelabrum cells are molecularly distinct, ubiquitous interneurons of the cerebellar cortex with specialized circuit properties. bioRxiv:2021.2004.2009.439172.

Osorno T, Rudolph S, Nguyen T, Kozareva V, Nadaf NM, Norton A, Macosko EZ, Lee WA, Regehr WG (2022) Candelabrum cells are ubiquitous cerebellar cortex interneurons with specialized circuit properties. Nature neuroscience 25:702–713.

Ozden I, Sullivan MR, Lee HM, Wang SS (2009) Reliable coding emerges from coactivation of climbing fibers in microbands of cerebellar Purkinje neurons. The Journal of neuroscience : the official journal of the Society for Neuroscience 29:10463–10473.

Pachitariu M, Steinmetz N, Kadir S, Carandini M, Kenneth D. H (2016) Kilosort: realtime spike-sorting for extracellular electrophysiology with hundreds of channels. bioRxiv:061481.

Paulsen O, Moser EI (1998) A model of hippocampal memory encoding and retrieval: GABAergic control of synaptic plasticity. Trends in neurosciences 21:273–278.

Piochon C, Kloth AD, Grasselli G, Titley HK, Nakayama H, Hashimoto K, Wan V, Simmons DH, Eissa T, Nakatani J, Cherskov A, Miyazaki T, Watanabe M, Takumi T, Kano M, Wang SS, Hansel C (2014) Cerebellar plasticity and motor learning deficits in a copy-number variation mouse model of autism. Nature communications 5:5586.

Rajasethupathy P, Sankaran S, Marshel JH, Kim CK, Ferenczi E, Lee SY, Berndt A, Ramakrishnan C, Jaffe A, Lo M, Liston C, Deisseroth K (2015) Projections from neocortex mediate top-down control of memory retrieval. Nature 526:653–659.

Raymond JL, Lisberger SG (1998) Neural learning rules for the vestibulo-ocular reflex. The Journal of neuroscience : the official journal of the Society for Neuroscience 18:9112–9129.

Raymond JL, Medina JF (2018) Computational Principles of Supervised Learning in the Cerebellum. Annual review of neuroscience 41:233–253.

Rieubland S, Roth A, Hausser M (2014) Structured connectivity in cerebellar inhibitory networks. Neuron 81:913–929.

Robinson DA (2022) Plasticity and repair of the vestibulo-ocular reflex. Prog Brain Res 267:183–214.

Rowan MJM, Bonnan A, Zhang K, Amat SB, Kikuchi C, Taniguchi H, Augustine GJ, Christie JM (2018) Graded Control of Climbing-Fiber-Mediated Plasticity and Learning by Inhibition in the Cerebellum. Neuron 99:999–1015.e1016.

Silva NT, Ramírez-Buriticá J, Pritchett DL, Carey MR (2022) Neural instructive signals for associative cerebellar learning. bioRxiv:2022.2004.2018.488634.

Simpson JI, Alley KE (1974) Visual climbing fiber input to rabbit vestibulo-cerebellum: a source of direction-specific information. Brain research 82:302–308.

Stahl JS (2004) Eye movements of the murine P/Q calcium channel mutant rocker, and the impact of aging. Journal of neurophysiology 91:2066–2078.

Stahl JS, van Alphen AM, De Zeeuw CI (2000) A comparison of video and magnetic search coil recordings of mouse eye movements. Journal of neuroscience methods 99:101–110.

Stone LS, Lisberger SG (1990) Visual responses of Purkinje cells in the cerebellar flocculus during smooth-pursuit eye movements in monkeys. II. Complex spikes. Journal of neurophysiology 63:1262–1275.

Tang X, Jaenisch R, Sur M (2021) The role of GABAergic signalling in neurodevelopmental disorders. Nature reviews Neuroscience 22:290–307.

Voges K, Wu B, Post L, Schonewille M, De Zeeuw CI (2017) Mechanisms underlying vestibulo-cerebellar motor learning in mice depend on movement direction. The Journal of physiology 595:5301–5326.

Witter L, Rudolph S, Pressler RT, Lahlaf SI, Regehr WG (2016) Purkinje Cell Collaterals Enable Output Signals from the Cerebellar Cortex to Feed Back to Purkinje Cells and Interneurons. Neuron 91:312–319.

