## Supplementary figures and images for "Molecular layer disinhibition unlocks climbing-fiber-instructed motor learning in the cerebellum"

### Supplemental Figure 1

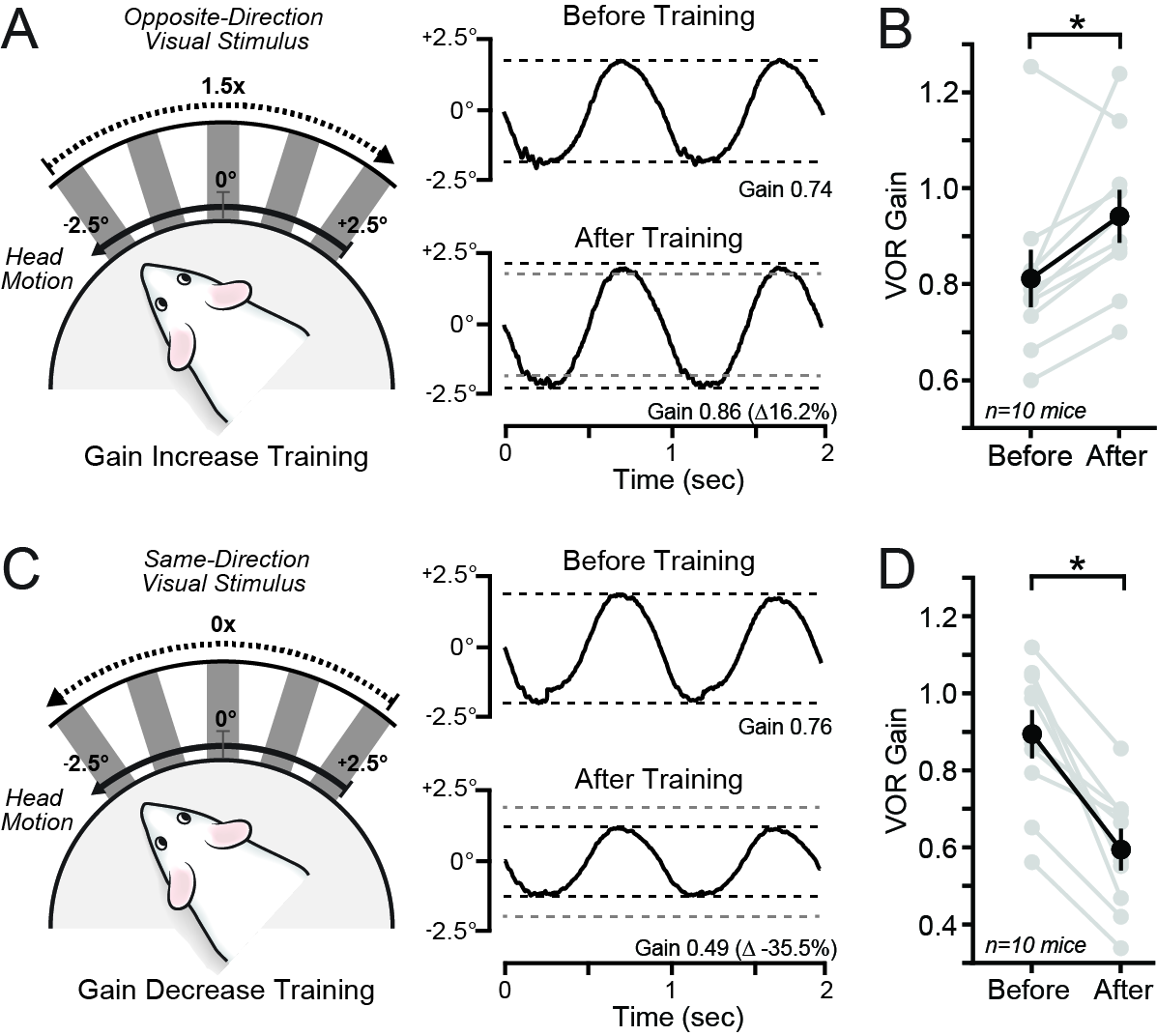

### Supplemental Figure 2

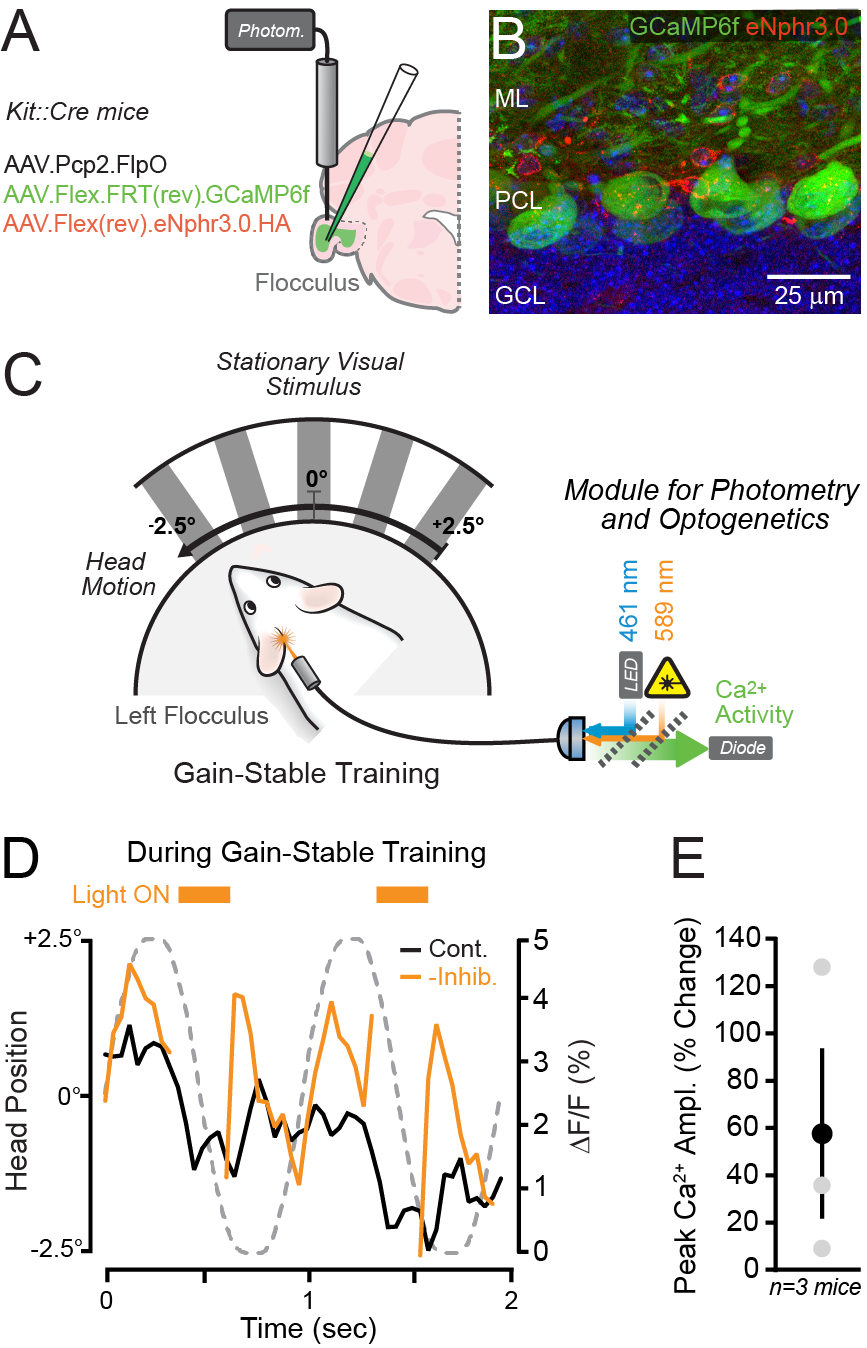
